# Single cell spatial proteomics maps human liver zonation patterns and their vulnerability to fibrosis

**DOI:** 10.1101/2025.04.13.648568

**Authors:** Caroline A. M. Weiss, Lauryn A. Brown, Lucas Miranda, Paolo Pellizzoni, Shani Ben-Moshe, Sophia Steigerwald, Kirsten Remmert, Jonathan Hernandez, Karsten Borgwardt, Florian A. Rosenberger, Natalie Porat-Shliom, Matthias Mann

## Abstract

Understanding protein distribution patterns across tissue architecture is crucial for deciphering organ function in health and disease. Here, we applied single-cell Deep Visual Proteomics to perform spatially-resolved proteome analysis of individual cells in native tissue. We combined this with a novel strategic cell selection pipeline and a continuous protein gradient mapping framework to investigate larger clinical cohorts. We generated a comprehensive spatial map of the human hepatic proteome by analyzing hundreds of individual hepatocytes from 18 individuals. Among more than 2,500 proteins per cell about half exhibited zonated expression patterns. Cross-species comparison with mouse data revealed conserved metabolic functions and human-specific features of liver zonation. Analysis of fibrotic samples demonstrated widespread disruption of protein zonation, with pericentral proteins being particularly susceptible. Our study provides a comprehensive resource of human liver organization while establishing a broadly applicable framework for spatial proteomics analyses along tissue gradients.

## Introduction

The liver, the largest internal organ, is crucial in orchestrating essential processes such as glucose homeostasis, protein synthesis, bile production, and detoxifying harmful substances. To efficiently manage these diverse functions, the liver has evolved a unique anatomical organization. Directional blood flow through the hepatic lobule, the smallest functional unit, creates distinct metabolic and signaling gradients along this axis. Although histologically hepatocytes appear identical, they perform very different and often opposing metabolic functions depending on their location within the lobule - a phenomenon known as liver zonation^1–4^. For instance, periportal hepatocytes primarily engage in gluconeogenesis and fatty acid oxidation, while their pericentral counterparts specialize in glycolysis and lipogenesis (**Fig. 1a**)^5^. This sophisticated spatial division of labor allows the liver to perform competing metabolic processes simultaneously with remarkable efficiency.

**Fig. 1.**
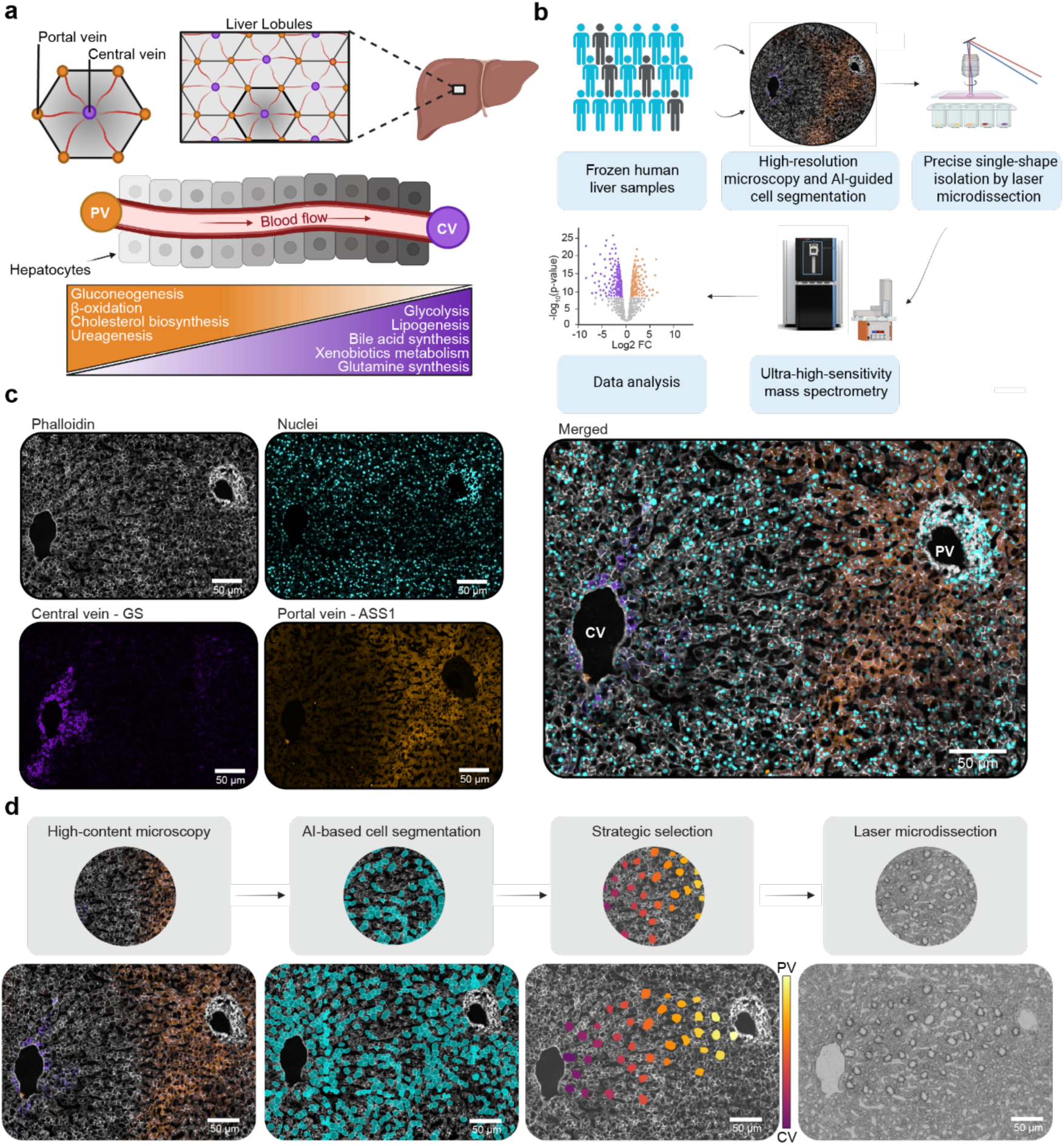
Single-cell Deep Visual Proteomics (scDVP) for spatially resolved proteomic analysis of liver zonation. **a)** Schematic representation of liver architecture showing the anatomical and functional organization of metabolic processes along the portal-central axis in human liver lobules. **b)** Workflow overview integrating high-resolution microscopy, AI-guided cell segmentation, laser microdissection, and ultra-high-sensitivity mass spectrometry for spatial proteomic analysis. **c)** Immunofluorescence imaging of human liver tissue visualizing glutamine synthetase (GS; pericentral), argininosuccinate synthetase 1 (ASS1; periportal), cell boundaries with phalloidin, and nuclei. **d)** Novel single-cell selection strategy of segmented contours (turquoise), strategic selection (color gradient), and laser microdissection of equally distributed cells along a zonation trajectory. PV: portal vein, CV: central vein. Scale bars, 50 μm.

Liver pathologies commonly develop and progress in a zonated manner, with different regions showing distinct susceptibilities to various disease processes^3,6^. Therefore, precise mapping of liver zonation, with enhanced spatial resolution, is essential for a thorough investigation of liver physiology and disease. Accurately quantifying proteins as functional players is particularly important in liver disease, where translational and post-translational regulation, including protein turnover, often drives pathological processes. While recent transcriptomic studies have advanced our understanding of zonation patterns^7–9^, transcripts do not always correlate with protein activities. Solid understanding at the protein level is vital for developing effective therapeutic strategies^10,11^.

Proteomic studies of liver zonation so far relied on histological techniques and immunostaining of selected proteins, potentially biasing investigations in already known directions. Mass spectrometry (MS)-based proteomics allows for a more in-depth and detailed analysis of tissue heterogeneity^12^. We have recently introduced Deep Visual Proteomics (DVP), a technology that integrates high-resolution microscopy with artificial intelligence (AI)-guided image analysis, automated laser microdissection, and ultra-sensitive MS^13^. DVP permits the analysis of collections of phenotypically similar cells from the same region of the intact tissue while maintaining vital spatial context. Recently, we further developed this technology to a single-cell proteomic workflow^13^, enabling protein expression analysis in individual cells within their native tissue environment^12,15,16^. We showcased this technology on mouse liver sections, and found precisely tuned metabolic activities along spatial gradients between the portal and central veins^15^. However, whether this spatial heterogeneity is conserved in the human liver remains unknown.

Here, we have substantially enhanced the technological framework of scDVP, enabling the precise quantification of proteins in hundreds of individual hepatocytes from a diverse cohort of 18 human liver samples. We developed an automated, strategic cell selection algorithm that streamlines the analysis of larger cohorts and maximizes information yield per tissue section. Furthermore, we have established a robust statistical framework to quantify protein expression gradients along spatial trajectories without artificial binning, making our approach applicable to a wide range of biological samples. These advancements allowed us to comprehensively characterize human liver tissues, creating the first single-cell proteomic map of protein gradients along the zonation axis in both healthy and disease states.

## Results

### Trajectory-driven cell selection enables scalable spatial proteomics in human liver

To create a comprehensive map of the proteome of human hepatocyte zonation at a single-cell resolution, we applied single-cell Deep Visual Proteomics (scDVP) to a cohort of healthy and fibrotic human liver tissue across 18 individuals (14 healthy and 4 fibrotic, **Fig. 1b**). The cohort included both female and male individuals from a broad range of ages and BMIs (see **Extended Data Table 1**). Images of the liver lobule sections were acquired using a four-channel immunofluorescence approach, to gain critical spatial context. In addition to cell borders stained by phalloidin and nuclei stained by DAPI which facilitate hepatocyte identification and segmentation, we also simultaneously stained for pericentral regions (glutamine synthetase, GS), and periportal regions (argininosuccinate synthetase 1, ASS1) (**Fig. 1c**). These inversely zonated hepatocyte landmark proteins allow spatial navigation and enable us to identify cell trajectories along the porto-central axis.

Our study presents a substantial increase in cohort size compared to previous scDVP efforts, which typically examined fewer than three individuals^15,17^. To facilitate the increased scale we aimed to maximize information yield from each tissue section while minimizing the number of samples to process. We developed a framework for automated, strategic cell selection enabling the comprehensive analysis of one single zonation trajectory per sample. The pipeline begins with an automated identification and characterization of central and portal veins based on the marker proteins GS and ASS1, respectively. After manual selection of a clear zonation trajectory between identified veins, our algorithm employs a combinatorial optimization approach called farthest-first traversal to select segmented hepatocytes which are distributed uniformly along the portal-to-central axis (see **Methods**)^18,19^. This approach maximizes the minimum distance between any two selected cells, ensuring comprehensive coverage of the zonation gradient while maintaining compatibility with downstream laser microdissection. Furthermore, comprehensive spatial metadata is recorded for each captured cell, including the critical spatial ratio *S* - a normalized measure ranging from 0 (central vein) to 1 (portal vein) that quantifies the relative position along the zonation axis (**Fig. 1d and Extended Data Fig. 1**). Forty-four cells per liver zonation trajectory were selected providing dense coverage of the approximately 20 hepatocytes that span a human liver lobule (**Extended Data Fig. 2**). By analyzing one optimal trajectory rather than dispersed cells, we capture the complete zonation gradient with fewer measurements while minimizing inter-lobule heterogeneity.

### Mapping the zonated proteome at over 2,500 proteins per single hepatocyte

We isolated a total of 792 single hepatocyte shapes with a median area of 651.4 μm², corresponding to the expected cross-section of human hepatocytes in 10 μm tissue slices (**Extended Data Fig. 3a**)^15^. Given the average hepatocyte diameter of 20 to 30 µm, and tissue sections of 10 µm thickness, microdissected samples corresponded to one-third to half a cell with approximately 250 pg of input material. For MS-analysis, we harnessed the recently introduced Orbitrap Astral mass spectrometer, which represents a significant advancement in sensitivity compared to previous instrumentation^12^. We designed a sample-tailored data independent acquisition (DIA) method which optimized the width of isolation windows based on the precursor density distribution of our samples (see **Extended Data Table 2**). This allowed efficient ion selection by employing narrower windows where precursor density is high and wider windows in sparse regions, maximizing peptide detection while minimizing interference. The resulting optimal coverage of existing precursors is especially critical when analyzing ultra-low input samples^20,21^.

With these advances, we identified a median of 2,539 proteins per shape and up to 3,926 proteins after applying quality threshold filtering (**Fig. 2a** and **Extended Data Fig. 3b**). We identified 5175 distinct proteins across our cohort with the known hepatocyte markers CPS1 and HGMCS2 among the highest abundant proteins^22,23^ (**Extended Data Fig. 3c**). This marks a strong improvement in protein quantification over our previous scDVP study in mice, with a median of 1,712 proteins per single shape. Moreover, in contrast to that previous effort the current study was measured in a label-free manner but yet at twice the throughput due to a much shorter Whisper gradients corresponding to 80 samples per day (SPD, see **Methods**). Total measuring time was only 10 days for this comprehensive single-cell proteomics study of 792 shapes. The LC-MS workflow showed high robustness across individuals, with a consistent median number of proteins identified and coefficients of variation, and absence of clustering of individuals, as reflected in principal components analysis (PCA, **Extended Data Fig. 3d-f**). Among samples from healthy individuals, we found a strong correlation between the position along the portal-central axis and the main component in PCA, indicating that space is the main driver of data variance (Spearman’s r = -0.55, pval = 3 ∗ 10^!”#^, n = 413, N = 14, **Fig. 2b)**.

**Fig. 2.**
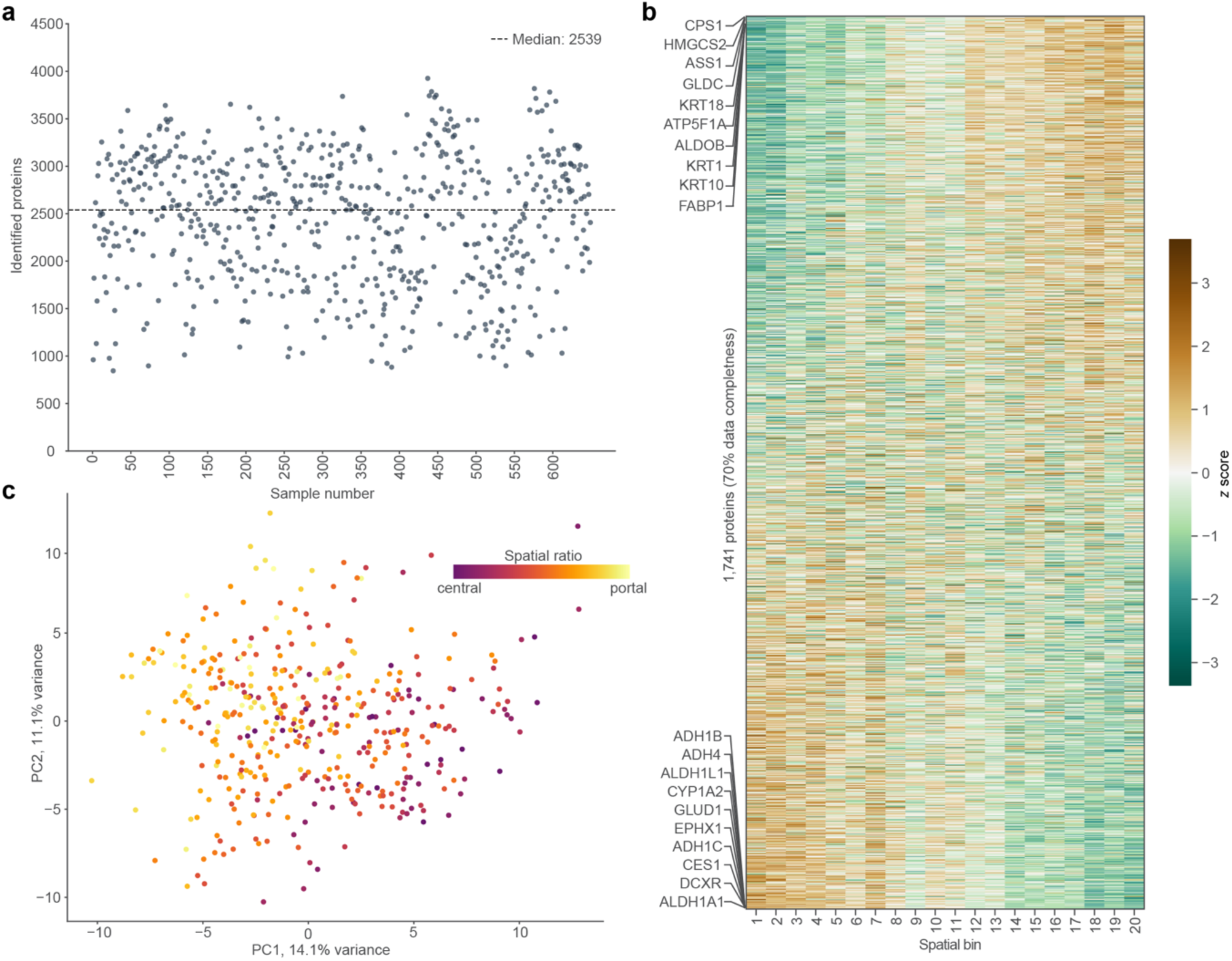
Deep proteome coverage enables spatial analysis of human liver zonation. **a)** Number of proteins identified per hepatocyte shape after quality filtering (n = 649, N = 18). The dotted line indicates the median depth across all included samples at 2,539 proteins. **b)** Principal component analysis of single-cell proteomes after outlier removal from healthy liver tissue. Each dot represents a single hepatocyte shape, with its color indicating spatial ratio *S* (n = 413, N = 14). **c)** Protein expression heatmap (*z* scored) from healthy individuals across 20 equal-width spatial bins from central (spatial ratio *S* = 0) to portal (spatial ratio *S* = 1). Proteins are ordered by expression differences from portal (top) to central vein (bottom). The top 10 proteins in each direction are labeled. Unless otherwise stated, only proteins detected in 70% of samples are included (n = 1741, N = 14).

Spatial analysis of the human liver proteome has so far relied on discrete binning approaches, commonly using three zones along the zonation axis^3^. More recent efforts increased the number of bins to 20 to minimize the averaging of effects^24^, we first sorted the hepatocytes into 20 equal-width bins along the trajectory according to their spatial ratio *S*, and then calculated the mean protein expression in each bin. Sorting the proteins by their expression change, clearly visualized the spatially dependent expression of the human liver proteome, known as liver zonation (**Fig. 2c**)^1–4^, consistent with previous findings in mice^15^. Hierarchical clustering of the binned data identified distinct protein expression clusters. Overrepresentation analysis of the two largest clusters revealed functional pathways characteristic for portal and central regions, such as oxidative phosphorylation or bile acid metabolism, respectively (**Extended Data Fig. 3g**). Together, our analysis demonstrates that the single-cell dataset captures the human hepatic proteome with high spatial fidelity.

### Single-cell protein gradient mapping defines spatial-dependent functionality of hepatocytes

Isolating single shapes by laser microdissection permits greatly increased spatial resolution. To leverage this aspect of scDVP and avoid reducing data content via binning, we developed a statistical framework that analyzes protein gradients continuously across space. Our approach is based on a linear mixed model, accounting for patient-specific variations, and includes proteins with a data completeness of at least 70% (see **Methods**)^25^. Using the spatial ratio *S* as an independent variable, we modeled protein expression values as the dependent variable. This determined the parameter β_1_, henceforth referred to as zonation coefficient, which characterizes and quantifies the protein expression gradient i.e., the rate at which the expression changes along the zonation trajectory (**Fig. 3a**). Furthermore, we calculated a statistical q-value by testing against the null hypothesis that gradients are zero. To further evaluate the continuous analysis approach, we performed a one-way ANOVA on the binned analysis and compared the sensitivity of the two methodologies. While both strategies show similar trends at higher q values with 429 proteins significant in both (Spearman correlation = 0.720), the continuous approach detected 397 statistically significant proteins that were not detected in ANOVA, whereas only 22 proteins were significant only in ANOVA (**Extended Data Fig. 4a**). This result clearly demonstrates the higher statistical power of the continuous analysis compared to ANOVA of binned lobule areas.

**Fig. 3.**
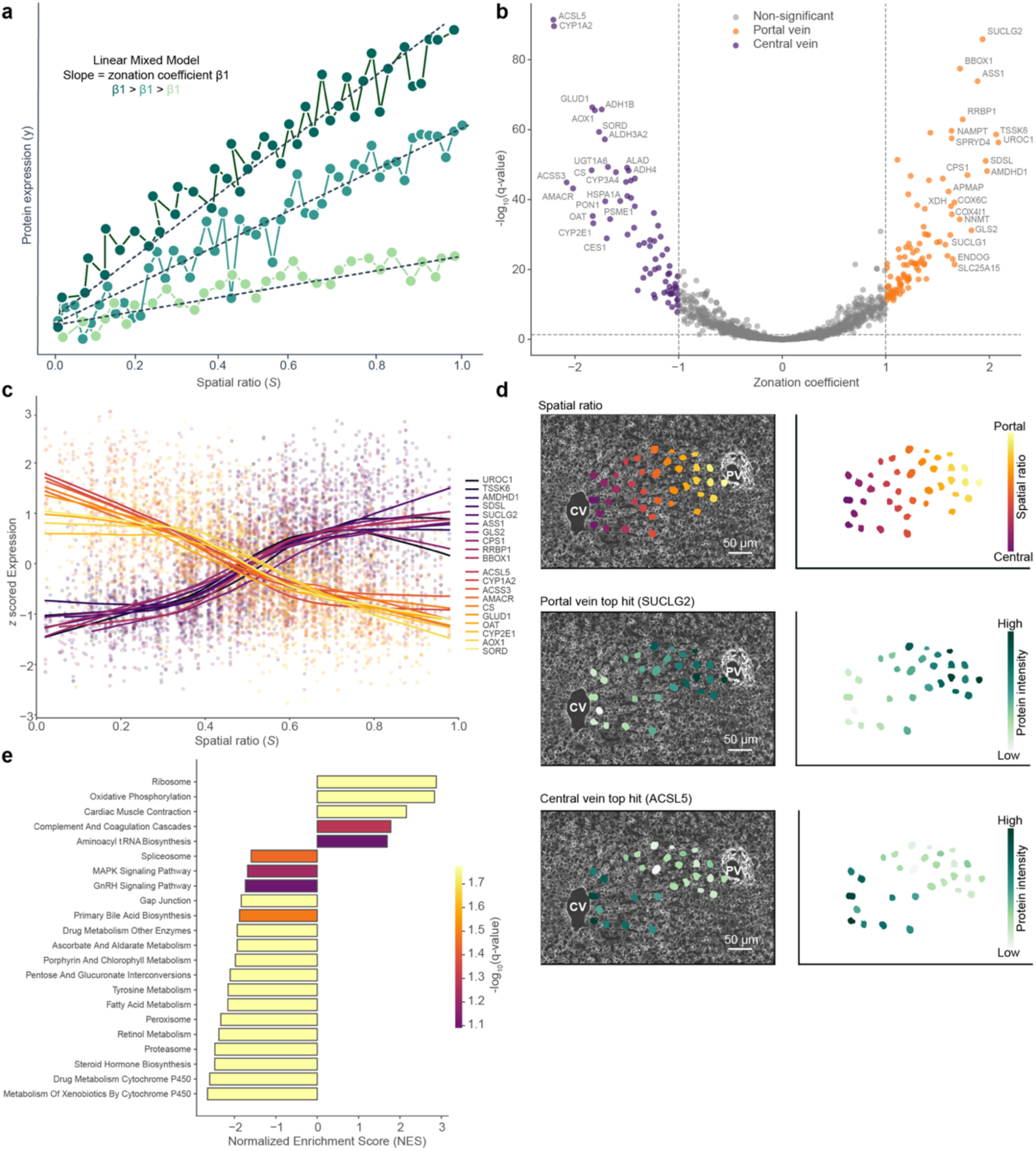
Gradient mapping of protein zonation patterns in healthy liver tissue. **a)** Schematic representation of continuous analysis approach based on a linear mixed model to quantify protein zonation patterns. Each line represents a protein expression pattern across the portal-central axis after normalization, with steeper slopes (β1, zonation coefficient) indicating stronger zonation. The model accounts for patient-specific variations through random intercepts. **b)** Volcano plot showing results of the continuous analysis. Each protein zonation coefficient (β1) determined by linear mixed modeling is plotted against the -log10(q-value) from a Wald test, which compares β1 against zero. The top 20 proteins from each side are labeled. Proteins with |zonation coefficient| > 1 and q < 0.05 were defined as strongly zonated. **c)** Expression profiles of the top 20 most significantly zonated proteins (purple shades: central vein, orange shades: portal vein). Individual measurements are shown as dots with LOWESS smoothing curves (fraction=0.4). **d)** Data visualization of protein zonation on microscopy image. Top: Example trajectory with spatial ratio color scheme. Middle and bottom: Protein intensities (green, low to high) of the most significantly zonated proteins in portal (SUCLG2) and central (ACSL5) regions. **e)** Gene set enrichment analysis ranked by zonation coefficient using KEGG pathway database. Normalized enrichment scores were shown with statistical significance assessed using Benjamini-Hochberg correction for multiple testing (color gradient, q < 0.1). (*continues on next page*) (*continuation from previous page*) Negative NES indicated enrichment towards the central vein, and positive NES enrichment towards the portal vein. Unless stated otherwise, only proteins with 70% data completeness are included (n = 1741, N = 14). PV: portal vein, CV: central vein. Scale bars, 50 μm.

Roughly in line with previous estimations, 47.4% of the identified proteins show a significantly zonated expression pattern (q < 0.05) in line with previous findings^7,10,15,26^. The observation that nearly half of the proteome within a morphologically identical cell type exhibits spatial organization underscores the remarkable single-cell diversity of hepatocytes and further highlights the importance of spatial context. By definition, negative coefficients of the spatial ratio S, indicate proteins enriched towards the central vein, while positive zonation coefficients represent proteins with higher expression towards the portal vein. Furthermore, their magnitude is a quantitative measure of zonation strength, enabling us to rank proteins by their degree of spatial organization. Stringent thresholding on absolute zonation coefficients > 1, identified 171 strongly zonated proteins (|zonation coefficient| > 1 and q < 0.05), with 69 showing central vein enrichment and 102 enriched towards the portal vein (**Fig. 3b**). A summary of the detected proteins together with an individual quantitative measure of zonation is provided as a resource for the community (see **Extended Data Table 3**).

We visualized the top 20 significantly zonated proteins at the cohort level, revealing clear and continuous gradients along the portal-central axis (**Fig. 3c**). This trend was consistently present for the two most significantly zonated proteins SUCLG2 and ACSL5, across individuals and when projected back onto the original microscopic image of one individual (**Fig. 3d** and **Extended Data Fig. 4b**). This ability to map MS-based proteomics data back onto the original microscopy images illustrates the unique strength of DVP to connect molecular profiles with spatial tissue context, bridging the gap between imaging and functional data as represented by proteome-wide analyses. Specifically, gene set enrichment analysis (GSEA) of proteins ranked by their zonation coefficient revealed distinct functional organization between portal and central regions in healthy liver tissue, indicated by positive and negative normalized enrichment scores (NES), respectively (**Fig. 3e**). Portal regions showed high enrichment of ribosomal and oxidative phosphorylation components. In contrast, central regions were characterized by elevated xenobiotic and steroid hormone metabolism. This spatial separation of key metabolic functions recapitulates known functional compartmentalization of the liver lobule, where periportal hepatocytes specialize in energy-intensive processes while pericentral hepatocytes focus on detoxification but provid rich additional molecular context^3,4^.

### Cross-species comparison reveals conserved and unique features of liver zonation

To investigate the extent of conservation of liver zonation between species, we compared this human dataset to our previously published mouse scDVP dataset^15^. Despite some methodological differences between the two studies, including different mass spectrometry platforms (Orbitrap Astral versus timsTOF SCP), sample preparation workflows (label-free versus dimethyl-labeled), and tissue types (surgically resected human versus perfused mouse liver), 2,829 homologous proteins where shared across this evolutionary distance of 75 million years^27^ (**Fig. 4a**). Applying our continuous gradient mapping framework on the mouse dataset identified 74 central and 94 portal vein-associated proteins with the stringent threshold filter (**Extended Data Fig. 5a**), consistent with the original study^15^. Zonation coefficients in human and mouse datasets indicated an overall similar spatial distribution of protein expression profiles (Pearson’s R = 0.44, **Fig. 4b**). Twenty-five central and 18 portal vein-specific proteins maintained their strong zonated profile across both species (**Fig. 4c**). These evolutionarily conserved zonated proteins include key enzymes involved in critical liver functions such as urea cycle components (ASS1, CPS1), xenobiotic metabolism (CYP2E1, CYP1A2), and energy production (NDUFS1, SUCLG1), suggesting their fundamental importance for liver physiology across mammals.

**Fig. 4.**
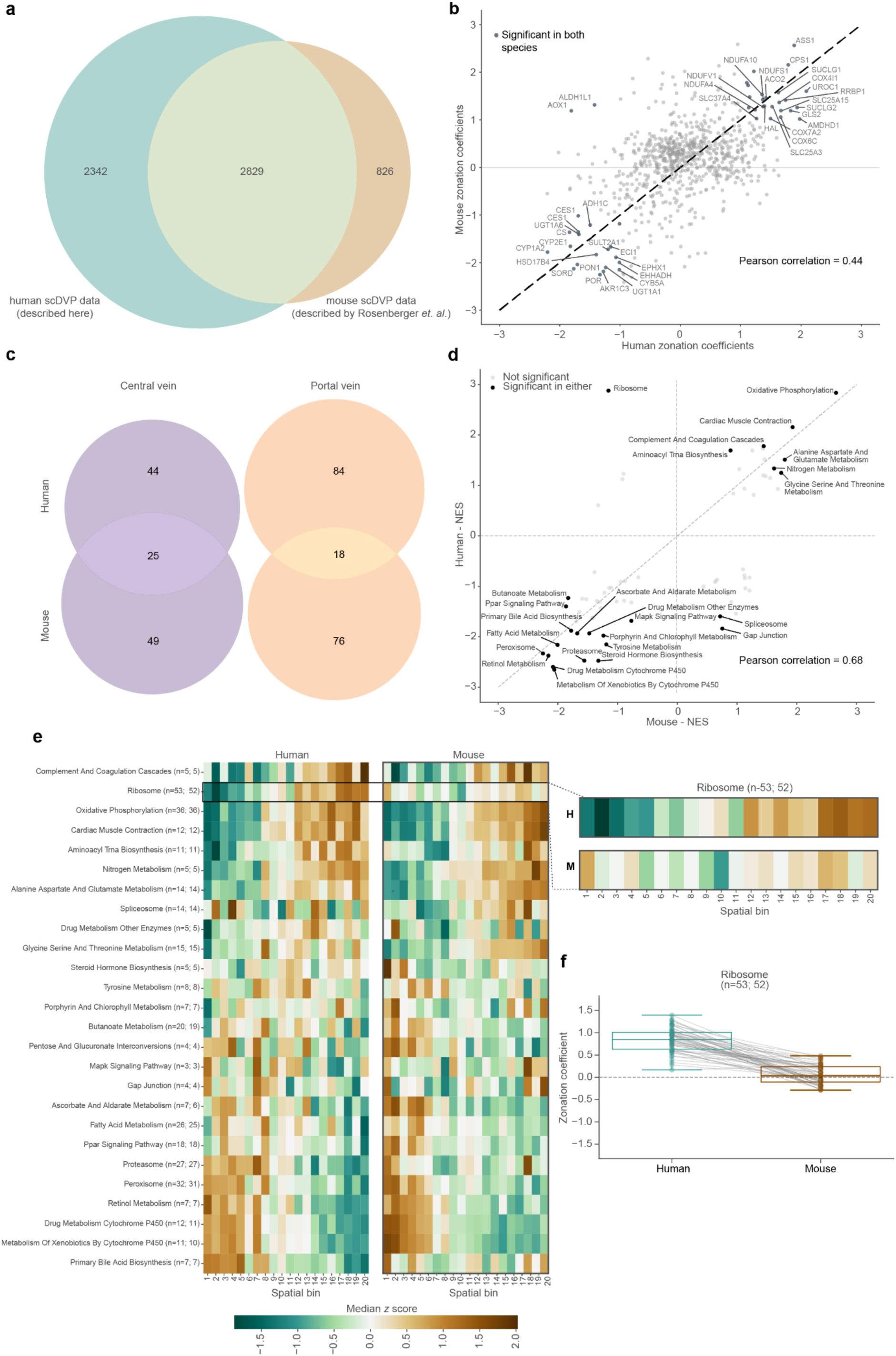
Cross-species comparison of liver zonation patterns between mouse and human. **a)** Overlap between proteins quantified in healthy humans (this study) and previously published mouse scDVP datasets (human: n = 5174, N = 14; mouse: n = 3655, N = 3) **b)** Comparison of zonation coefficients determined by continuous analysis between humans and mouse for shared proteins. Proteins with a |zonation coefficient| > 1 in both species are labeled. **c)** Comparison of differentially expressed portal and central vein proteins in mice and humans. **d)** Comparison of normalized enrichment scores (NES) from independent gene set enrichment analysis (KEGG pathways) of human and mouse datasets, ranked by the zonation coefficient. Statistical significance is assessed using Benjamini-Hochberg correction for multiple testing, and significantly enriched pathways (q < 0.1) in either species are labeled. **e)** Heatmaps comparing expression patterns (*z* scored) of shared proteins across 20 equally-spaced spatial bins from central (spatial ratio *S* = 0) to portal vein (spatial ratio *S* = 1) for significantly enriched pathways (q < 0.1). The median *z* score of the proteins assigned to the respective pathway in each bin is displayed. The right panel shows a closeup of the ribosomal protein expression patterns in humans (H) and mice (M). Numbers indicate the number of proteinsin the respective term (H; M). **f)** Comparison of ribosomal proteins zonation coefficients measured in human and mouse datasets (boxes show first and third quartiles, center line indicates median, whiskers extend to 1.5x interquartile range). Individual proteins are shown as dots with connecting lines between matched proteins. Colors indicate human (turquoise) and mouse (brown) datasets. Unless stated otherwise, only proteins at 70% data completeness are included (human: n = 1741, N = 14; mouse: n = 981, N = 3).

Of note, many proteins were zonated in one species but not the other, including two ambiguous homologous pairs (**Extended Data Fig. 5c-d**). We performed GSEA independently for each species for insights at the pathway level, ranking proteins by their zonation coefficients. The resulting NES values were highly similar between species, with a stronger correlation at the pathway level than at the individual protein level, reflecting greater conservation of functional modules than their individual components (Pearson’s R = 0.68, **Fig. 4d**). Visualizing protein zonation coefficients aggregated by functional pathways (q < 0.1), revealed remarkably similar spatial patterns between mouse and human, exemplified by oxidative phosphorylation enrichment in the portal vein and enrichment of xenobiotic metabolism in the central vein (**Fig. 4e**). Interestingly, the zonation pattern of the abundant group of ribosomal proteins strikingly diverged patterns between species. In humans, all detected ribosomal proteins consistently showed clear portal vein enrichment - a pattern that was absent in mice at both pathway and individual protein levels (**Fig. 4e-f**, **Extended Data Fig. 5d**). This difference suggests a species-specific regulation of protein synthesis capacity across the liver lobule. The human-specific periportal enrichment of ribosomal proteins may reflect heightened protein synthesis demands in oxygen-rich regions to support the production of plasma proteins and other secreted factors.

### Liver fibrosis asymmetrically disrupts protein zonation

To investigate how liver disease affects spatial protein organization, we analyzed liver sections from patients with fibrosis. Fibrotic status was assessed histologically by Masson Trichrome staining performed by a single pathologist. Liver samples with collagen-positive areas exceeding 5% of the tissue were classified as fibrotic. This analysis identified 14 tissues as healthy and 4 as fibrotic in our cohort (**Extended Data Fig. 6a-b**). To explore the spatial heterogeneity of histologically functional hepatocytes, we selected trajectories from regions that appeared morphologically normal but were adjacent to fibrotic areas (**Extended Data Fig. 2**). The fibrotic samples were analyzed in a similar manner to their healthy counterparts.

Comparing protein zonation coefficients between healthy and fibrotic tissue revealed a general loss of spatial organization. This was demonstrated by zonation coefficients shifting closer to zero for both periportal and pericentral zonated proteins, when compared to their corresponding proteins in healthy individuals. We quantified this systematic trend by measuring the deviation of the observed regression (solid line) from the expected diagonal of unchanged coefficients (dashed line), with the flatter slope indicating global reduction of zonation magnitude in fibrosis (**Fig. 5a)**. Notably, while proteins showed reduced zonation, we did not observe any inversion of zonation patterns between conditions (**Fig. 5a**). PCA of the fibrotic samples showed a weaker association between the spatial ratio and main component than in the healthy cohort (Spearman’s r = -0.31, pval = 1.7 ∗ 10^!”^, **Extended Data Fig. 6c**).

**Fig. 5.**
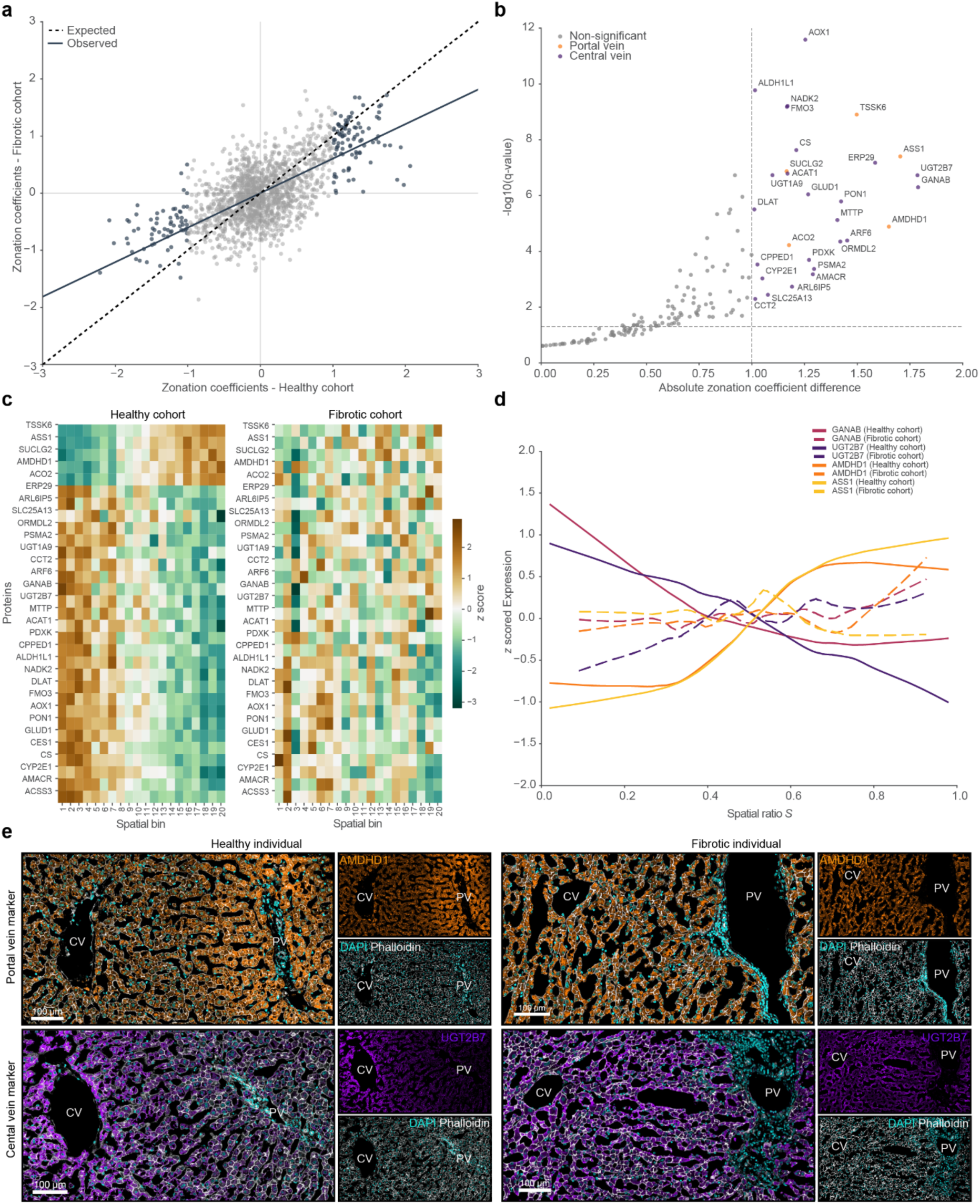
Loss of protein zonation patterns in human fibrotic liver. **a)** Zonation coefficients in healthy and fibrotic liver samples. Dotted line indicates theoretical unchanged zonation coefficients, solid line shows observed relationship as a linear regression line. **b)** Absolute differences in zonation coefficients between healthy and fibrotic tissue. Only strongly zonated proteins (|zonation coefficient| > 1 and q < 0.05) from healthy are analyzed (n = 171). Proteins with significant zonation loss are highlighted and color-coded by their original spatial enrichment. Their zonation coefficients were compared by Wald test (absolute zonation coefficient difference > 1 and q < 0.05). **c)** Protein expression heatmap (*z* scored) from significant proteins in **c)** in the healthy and fibrotic cohort. 20 equal-width spatial bins from central (spatial ratio *S* = 0) to portal (spatial ratio *S* = 1) are shown. **d)** Expression profiles of the four proteins with the strongest absolute zonation coefficient difference. (*continues on next page*) (*continuation from previous page*) LOWESS smoothing curves (fraction=0.4) are shown, with solid lines indicating the healthy and dotted lines fibrotic cohort. **e)** Immunofluorescence staining of a portal vein (top) and central vein (bottom) associated protein in healthy (left; individual 12) versus fibrotic (right; individual 16) tissue. Unless otherwise stated, only proteins at 70% data completeness are included (healthy: n = 1741, N = 14; fibrosis: n = 1684, N = 4). PV: portal vein, CV: central vein. Scale bars, 100 μm.

To statistically evaluate the proteins driving the reduction in the global zonation profile we compared the observed difference in zonation coefficients (Δβ_1_) to the null hypothesis of a zero difference. This analysis identified a subset of proteins (18.1%) with particularly dramatic reductions in their zonation coefficients in fibrotic tissue, including key metabolic enzymes from both portal vein-associated pathways, such as ASS1, SUCLG2 and AMDHD1, as well as central vein processes, exemplified by CYP2E1, UGT2B7 and ALDH1L1 (**Fig. 5b** and **5c**). Interestingly, central vein-associated proteins were generally more strongly impacted in their spatial organization than their portal vein counterparts. We further highlighted proteins showing the strongest loss in their zonal expression distribution in the fibrotic samples by displaying the measured intensity distribution (**Fig. 5d)**. To validate our findings using an independent technology, we stained tissue samples from the same patients with immunofluorescence antibodies targeting proteins that showed a loss of zonation in our MS-based analysis. Immunofluorescence staining of AMDHD1 at the portal vein, as well as UGT2B7 at the central vein, confirmed the observed change in spatial organization (**Fig. 5e**).

As already suggested by the list of proteins with the strongest impact of fibrosis on their spatial organization, we found that the loss of protein zonation was generally not uniform across the liver lobule. Proteins that were enriched in healthy tissue near central vein regions showed a significantly stronger reduction in their zonation coefficients compared to portal vein proteins, with median zonation coefficient changes of Δ = 0.71 and Δ = 0.31, respectively (**Fig. 6a**). This discrepancy was also evident at the individual protein level. Overall, 41.2% of portal vein proteins preserved their zonation in fibrotic liver tissue, in contrast to only 11.6% of central vein proteins (**Fig. 6b**).

**Fig. 6.**
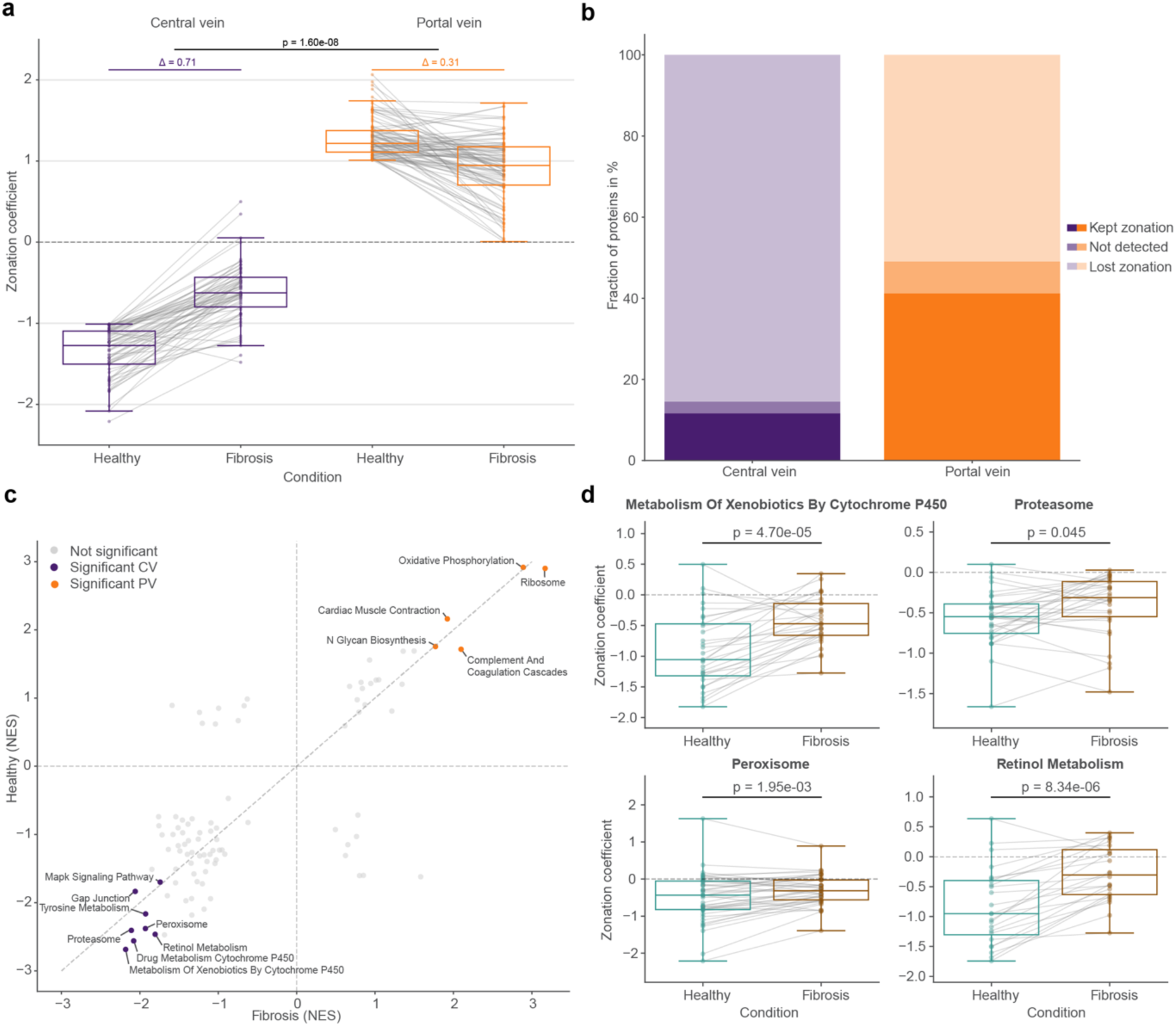
Fibrosis predominantly affects central vein protein zonation patterns. **a)** Zonation coefficients comparing healthy and fibrotic tissue. Strongly zonated proteins (|zonation coefficient| > 1 and q < 0.05) from the healthy cohort that are also detected in the fibrotic cohort are taken into account (central vein (CV): n = 67 and portal vein (PV) n = 94). Gray lines connect matched proteins between conditions. Delta values indicate the median absolute zonation coefficient change between conditions at the respective vein. The *p-*value is determined by a two-sided, unpaired rank sum test for the absolute differences between healthy and fibrosis for each protein. **b)** Fraction of proteins in central and portal vein showing lost (|zonation coefficient| < 1) or maintained zonation (|zonation coefficient| > 1 and q < 0.05), or not detected in fibrotic tissue. **c)** Correlation of normalized enrichment scores (NES) from independent GSEA analysis (KEGG pathways) of the healthy and fibrotic cohort, ranked by the zonation coefficient. Pathways significantly enriched (q < 0.1) in both cohorts are color coded. **d)** Box plots comparing zonation coefficients between healthy (turquoise) and fibrotic (brown) tissue for four selected central vein-enriched KEGG pathways. The *p* values are determined by a two-sided Wilcoxon test for paired samples. Boxes show first and third quartiles, center line indicates the median, whiskers extend to 1.5x interquartile range, and individual proteins are shown as dots with gray lines connecting matched proteins between conditions. Unless otherwise stated, only proteins at 70% data completeness are included (healthy: n = 1741, N = 14; fibrosis: n = 1684, N = 4).

Similar to our GSEA on zonation coefficients for healthy tissue, we performed this analysis for the fibrotic cohort (**Fig. 3e** and **Extended Data Fig. 6d**). The enriched terms remained consistent between the healthy and the fibrotic cohorts. However, comparing NES values between healthy and fibrotic tissue for pathways significant in both conditions corroborated the asymmetric change in spatial organization. Central vein-associated metabolic pathways particularly deviated from the diagonal line towards lower enrichment scores in fibrosis (**Fig. 6c**), as exemplified by several key metabolic pathways located at the central vein area, including xenobiotic metabolism, proteasome function, peroxisome pathways, and retinol metabolism. All of these processes moved towards a median zonation coefficient of zero, indicating a general loss of spatial organization in fibrotic tissue (**Fig. 6d**). Together, these results demonstrate that liver fibrosis leads to a pronounced disruption of protein zonation profiles, with central vein-specific processes being particularly vulnerable to spatial disorganization.

## Discussion

In this study, we advanced the scDVP workflow to enable the analysis of small clinical cohorts, creating an in-depth spatial proteome map of human liver tissue at single-cell resolution. Using a label-free MS approach, we quantified as many as 4,000 proteins per individual hepatocyte, achieving a median depth of 2,539 proteins across hundreds of cells (**Fig. 2a**). We developed an automated, strategic cell selection approach to identify cells with optimal distribution along a given spatial trajectory. This enabled efficient sampling of cells, maximizing information gain from each tissue section, thereby requiring fewer cells to capture the spatial organization of tissues. This represents an important technological advance for liver research and spatial proteomics.

Furthermore, we created a statistical framework to analyze protein expression changes continuously along spatial trajectories. The human liver is particularly suited to benchmark such an approach, as well-studied metabolic and signaling gradients occur over a relatively short physical distance of approximately 20 hepatocytes or about 385 µm in humans^28^. While earlier liver zonation studies relied on a three-zone model (periportal, mid-lobular, pericentral) or discretized the portal-to-central axis into twenty bins, the framework presented here captures the gradual nature of biological processes in the lobule without artificial boundaries^3,15^. Our approach models protein expression as continuous trajectories through space using linear mixed models, which account for patient-specific variations while avoiding the need to bin the spatial trajectory and reducing sensitivity to missing values, e.g. due to dropout samples (**Fig. 3a** and **Extended Data Fig. 4a**). This approach generates a quantitative measure of the degree of zonation for each protein. Our two-tiered statistical approach, combining q-values and zonation coefficient thresholds, enabled the identification of both subtle and strong zonation patterns.

In our study, we confirmed and extended previous findings indicating that half of the hepatic proteome is expressed with spatial variation, and171 proteins were found to be associated with a strong zonation profile here^7,10,15,26^. We characterized the functional zonation created by this extensive shift in the hepatic proteome along the porto-central axis, which creates an elegant molecular division of labor that optimizes overall tissue efficiency.The in-depth dataset of the human heapatic proteomes ranked by zonation behavior serves as a resource for researchers investigating liver physiology, disease mechanisms, and potential therapeutic targets within the spatial context in humans. While we demonstrated this framework on liver tissue, it is broadly applicable to any spatial proteomics dataset where gradual changes in protein expression occur, such as developmental gradients or tumor microenvironments.

We further re-analyzed our previously published mouse scDVP dataset^15^ using the continuous analysis framework introduced here and compared it with the human dataset (**Fig. 4**). While a relatively low number of zonated proteins overlapped between species, the spatial organization of core metabolic functions showed high conservation, indicating that liver zonation is remarkably conserved from mice to humans. Our results also highlight notable species-specific differences, most strikingly the portal zonation of ribosomal proteins that we observed only in humans (**Fig. 4e-f**). This suggests that while the spatial separation of incompatible processes is conserved, the regulation of zonal expression may vary across organisms. This finding may be useful when considering when to use mice to model human disease.

To examine the impact of pathological perturbation on zonation, we expanded our analysis to fibrotic human samples. Fibrosis disrupts lobular architecture and obstructs blood flow, with consequent changes in the oxygen gradient^29^. Pericentral hepatocytes, naturally exposed to lower oxygen levels, are profoundly influenced by this environment, as it regulates their gene expression through HIF1α^30^. At the proteome level, we found that fibrosis results in a marked disruption of protein zonation patterns, characterized by reduced expression of zonated proteins (**Fig. 5b**). Remarkably, fibrosis disproportionately impacted pericentral hepatocytes and related pathways, revealing their vulnerability to spatial disorganization (**Fig 6**). Current and future research will address the role of pericentral hepatocytes in adaptive responses^31^. Overall, these findings underscore the significance of fibrosis in altering the liver’s microenvironment and protein expression landscape.

While our gradient mapping analysis provides valuable insights into protein distribution patterns, it is primarily designed to identify linear protein expression gradients. Although non-linear gradients can still be captured, our model might not optimally fit these patterns. Additionally, the current state-of-the-art MS setup, while robust for single hepatocyte shapes containing about 250pg of protein (equivalent to one complete HeLa cell), will benefit from further technological advances in sensitivity to expand the application of scDVP to smaller cell types^24^.

In conclusion, combining automated cell selection and gradient mapping creates a powerful platform for spatial proteomics that can be readily adapted to diverse tissue samples and biological questions^12^. The in-depth spatial proteome atlas we generated provides a community resource for understanding human liver organization and may serve as a blueprint for studying protein gradients in other biological systems, from development to disease progression.

## Author Contributions

C.W., F.R., N.P.S., and M.M. conceptualized the study. C.W. and L.B. performed experimental work. C.W. and S.S. acquired MS data. C.W., L.B., L.M., P.P., S.B.M. and F.R. performed data analysis. L.M. and P.P. equally contributed to developing algorithms. J.H. and K.R. provided clinical samples and input. C.W. curated data. K.B., F.R, N.P.S. and M.M. supervised the project. C.W., F.R., N.P.S., and M.M. wrote the original manuscript draft. All authors read, revised and approved the manuscript.

## Acknowledgements

We thank our colleagues at the Department of Proteomics and Signal Transduction at the Max Planck Institute of Biochemistry, as well as our colleagues at the Center for Proteome Research in Copenhagen, for their input and support. We are particularly grateful for input and help from Marvin Thielert and Tim Heymann. F.A.R. is an EMBO postdoctoral fellow (ALTF 399-2021). This study has been supported by the Max Planck Society for Advancement of Science (M.M). The funders had no role in study design, data collection and analysis, the decision to publish, or the preparation of the manuscript. We likewise thank members of the Porat-Shliom lab for discussions and feedback, especially Dr. Kathrine Barrows, for guidance with the clinical database. This work was supported by the Intramural Research Program at the NIH, National Cancer Institute grant number 1ZIABC011828 (N.P.S).

## Competing Interests

M.M. is an indirect investor in Evosep. K.B. is co-founder and scientific advisor of Computomics GmbH, Tübingen, Germany. All other authors declare no competing interests.

## Methods

### Human tissue collection

Human tissue was acquired with informed consent under an NIH IRB-approved protocol (NCT01915225) for surgical resection or risk-reducing surgery performed on patients with germline CDH1 mutation(s). The tissue procured was characterized by either a normal or tumor-normal margin based upon visible inspection and confirmation of the final histopathologic examination. Following tissue procurement, which was completed roughly within 20 minutes of incision, tissue was fixed in 1% PFA in PBS for 48 or 72 hours, washed in PBS and incubated in 30% sucrose in PBS for at least 24 hours before embedding in OCT. OCT blocks were stored at -80 °C.

### Cryosectioning, tissue mounting and immunofluorescence

Two micrometer polyethylene naphthalate (PEN) membrane slides (MicroDissect GmbH) were pre-treated by UV exposure at 254 nm for 60 minutes to improve tissue adherence. Directly following UV treatment, sequential washing steps were performed on the slides: first in 350 mL acetone, then in a solution of 7 mL VECTABOND reagent (Vector Laboratories; SP-1800-7) diluted in 350 mL acetone, followed by a brief 30-second rinse in ddH_2_O. Slides were then dried under a gentle stream of nitrogen gas. 10 µm thick liver sections were thawed once and cut onto pre-cooled glass or PEN membrane slides using a Leica cryostat (Leica CM1950) for histology or Deep Visual Proteomics purposes, respectively. The slides were stored at -80°C.

For immunofluorescent staining, the slides were thawed at room temperature followed by a 2-minute fixation with 4% PFA in PBS at 37°C. The tissue was then permeabilized using 95% ethanol in ddH_2_O for 2 minutes at room temperature, rehydrated with PBS, and blocked with 5% BSA in PBS for at least 15 minutes at room temperature. Following this, sections were incubated with primary antibody Glutamine Synthetase (GS; abcam; ab176562; 1:300) overnight at 4 °C. The next day, the sections were washed, three times for 15 minutes each, with PBS and then incubated with the secondary antibody and fluorescent dyes Phalloidin (Thermo Fisher; A30104; 1:300), and Sytox Green (Thermo Fisher; S7020; 1:500) for 1 hour at room temperature. The following day, the sections were washed, three times for 15 minutes each, with PBS and then incubated at 4 °C overnight with primary antibody ASS1 (abcam; ab170952; 1:100) conjugated using Zenon™ Rabbit IgG Labeling Kits (Thermo Fisher; Z25307; 1:20 relative to microliters of ASS1 used). The next day, after an additional three washes, for 15 minutes each with PBS, the slides were left unmounted and stored at 4 °C until shipment.

A similar immunofluorescence protocol was used to validate protein distribution. Primary antibodies UGT2B7 (ProteinTech; 16661-1-AP; 1:50) or AMDHD1 (OriGene; TA809954; 1:250) were incubated overnight at 4 °C. The next day, the sections were washed, three times for 15 min each, with PBS and then incubated with secondary antibodies (1:500), and fluorescent dyes DAPI (Thermo Fisher Scientfic; H3570; 1:1000) and Phalloidin (Thermo Fisher Scientific; A22287; 1:300) for 1 hour at room temperature. After an additional three washes, for 15 min each with PBS, the slides were mounted in the dark overnight in preparation for imaging.

### Histology staining and fibrosis analysis

H&E, Oil Red O, and Masson Trichrome histological staining were performed on liver sections for pathological assessment using standard protocols by the Molecular Pathology Unit NCI. The slides were scanned using an Aperio AT2 scanner (Leica Biosystems, Buffalo Grove, IL) at 20X magnification, generating whole-slide digital images. Image analysis was conducted using HALO imaging software (version 3.6.4134.362; Indica Labs, Corrales, NM), with image annotations carried out by a single pathologist. The analysis was performed utilizing Area Quantification (version 2.1.11) for Masson’s trichrome in HALO to determine the percentage of positive area. Only tissue areas (excluding vessel lumen and white spaces) were taken into consideration by the software. Additionally, regions exhibiting artifacts, such as folds and tears as well as the liver capsule were excluded from the analysis.

### High-resolution microscopy and image processing

For scDVP, confocal image acquisition was performed using a PerkinElmer OperaPhenix high-content imaging system equipped with a 20× air objective (numerical aperture 0.8). The system was operated through Harmony software version 4.9, with images collected using 2×2 binning and 10% overlap between adjacent tiles. For each sample, a single focal plane was manually selected and maintained across all channels. To eliminate channel crosstalk, fluorophores were excited sequentially, with the following acquisition parameters: CFP at 80% laser power and 100ms exposure, Alexa 488 at 30% power and 20ms exposure, Alexa 568 at 80% power for 40ms, and Alexa 647 at 60% power for 40ms. Post-acquisition flat-field correction was applied through the Harmony v4.9 software platform. Image reconstruction was accomplished using scPortrait^32^, with the phalloidin-CFP signal used as the reference channel for calculating tile positions. These positions were subsequently applied to all channels to generate one tif-file per channel per sample. Stitching parameters were set to 0.1 for tile overlap, with filter sigma and maximum shift values of 1 and 50, respectively.

Immunofluorescence images for validation of loss of zonation profiles were acquired using a Leica SP8 inverted confocal laser scanning microscope, a 63x (1.4 NA) oil objective, a zoom of 1.00, and a pixel resolution of 1024×1024. The tile scans were merged using the Leica software and further processed using Imaris (Bitplane 9.9.0) and Image J.

### Identification and strategic selection of single cells

The processed and stitched images were imported into the Biological Image Analysis Software (BIAS) using the packaged import tool and retiled to 1024 x 1024 pixels with 10% overlap. The region of interest in each image was selected. Cell segmentation was performed externally using a custom-trained Cellpose segmentation model (Cellpose version 2.0) on the phalloidin staining channel^33^. The resulting masks were imported into BIAS where duplicates were removed. Three characteristic reference points were selected based on prominent tissue features per image. Contours and reference points were exported as .xml files. The contours were subsequently simplified by removing at least 90% of the data points defining the contour’s shape.

In this study, only cells belonging to one defined zonation trajectory per patient were analyzed. These were selected in a automated fashion: First, portal and central veins were automatically detected in the images using a custom algorithm by which areas with no cells were identified. A Gaussian blur and a morphological dilation was applied to the image to remove small voids and subsequently was thresholded using the filters.threshold_minimum function from the skimage package. Central and portal veins were then identified by the expression of the marker proteins glutamine synthase (GS) and argininosuccinate synthase (ASS1), respectively. All encountered veins were grouped by the algorithm in non-overlapping trajectories, defined by a pair of portal and central veins. The trajectory of interest was selected manually. Defined regions of interest were automatically generated, including all cells segmented directly between the veins and up to an angle of 65 degrees within the axis defined by the center of both veins (Fig. S1). Cells to segment were then chosen such that the minimum distance between any two shapes was maximized by solving a combinatorial optimization problem known as *remote-edge* diversity maximization using the farthest-first traversal algorithm^18,19^. Up to 44 non-overlapping shapes per pair of central and portal veins were selected to be cut via laser microdissection for further processing. For each of the selected cells, the spatial ratio S was calculated, as 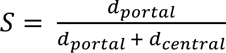, with *d_portal_* the distance (e.g., in pixels) from the center of the shape at hand to the portal vein and *d_central_* the distance to the central vein. The values from 0 (i.e., close to the central vein) to 1 (i.e., close to the portal vein) are covered uniformly by the selected shapes. To facilitate this analysis workflow, we developed a graphical user interface (GUI) where users only need to select two points of interest - point A (central vein) and point B (portal vein). An XML file containing the coordinates of the 44 cells along that trajectory is generated automatically by the GUI, enabling straightforward downstream analysis. Generally, only trajectories between clearly identified central and portal veins were analyzed.

### Laser microdissection

Following image alignment using three manually selected tissue reference points, cell contours were imported and isolated using a Leica LMD7. The system was operated in a semi-automated manner controlled by the LMD v8.3 software and using the following optimized parameters: 63x objective, power between 51-56, aperture 1, speed 10, middle pulse count 1, final pulse 0, offset 100, head current between 50-56%, and pulse frequency set to 2,900. The system was aligned to each plate as well as calibrated to account for gravitational stage shift when collecting into low-binding 384-well plates (Eppendorf 0030129547). Single cells were sorted in every other well and the outer rows and columns were left empty. A protective shield plate was positioned above the sample stage to avoid collection errors. The collection plates were sealed, centrifuged at 1,000g for 2 minutes and frozen at -20°C until further processing.

### Sample preparation and peptide loading

The following sample preparation steps were performed on an Agilent Bravo automated liquid handling platform in a semi-automated manner to minimize sample loss. Additionally, the plates were sealed with PCR sealing foil during incubation steps. Plates containing single cells were removed from -20°C and immediately centrifuged at 2000g for 2 minutes. The wells were washed with 28 μL 100% ACN and subsequent dried in a vacuum concentrator at 45°C for 20 minutes to ensure the presence of cut shapes at the bottom of the well. 6 μL lysis buffer (0.013% DDM in 60 mM TEAB buffer, pH 8.5) was added per well. Lysis was performed at 95°C for 60 minutes in PCR cycler (lid temperature 110°C). 1 μL of 80% ACN was added and the samples were cooked at 75°C for additional 60 minutes. After a brief cooling, 1 μL digestion mixture containing 4 ng/μL LysC and 6 ng/μL trypsin in 60 mM TEAB buffer was added and incubated overnight (approx. 18 hours) at 37°C. On the next day, the samples were immediately frozen at -20°C until sample loading was performed.

The samples were loaded on C-18 tips (Evotip Pure, EvoSep). The plates were thawed, immediately centrifuged at 2000g for 2 minutes and kept on ice until loading. EvoTips were activated in 1-propanol for 3 minutes. The tips were washed twice with 50 μL buffer B (0.1% formic acid, 99.9% ACN) and centrifuged at 700g for 1 minute between washes. A second activation in 1-propanol was performed for 3 minutes, followed by two washing steps with 50 μL buffer A (0.1% formic acid). The disk was kept wet after the final wash. 70 μL buffer A was added per tip and the samples were loaded. Each well was rinsed with 10 μL buffer A and added to the respective tip. For the peptide binding step, centrifugation was performed at 700g for 2 minutes. The tips were washed with 50 μL buffer A. Finally, 150 μL buffer A was added as overlay and the tips centrifuged at 700g for 15 seconds. The loaded tips were stored for maximum 3 days in an EvoTip box containing fresh buffer A until loading on the LC-MS.

### LC-MS/MS analysis

Analysis was performed on an Evosep One liquid chromatography system (Evosep) coupled to an Orbitrap Astral mass spectrometer (Thermo Fisher Scientific). An EASY-Spray source (Thermo Fisher Scientific) operating at 1900 V connected the two systems. Peptide separation was achieved using a Whisper Zoom 80 SPD gradient on an Aurora Rapid C18 column (5 cm, 75 μm ID, 1.7 μm particle size, IonOptics) at 60 °C. A throughput of 80 samples per day was achieved.

The Orbitrap Astral was equipped with a FAIMS Pro interface (-40 V compensation voltage, 3.5 L/min carrier gas, Thermo Fisher Scientific) and sample acquisition was exclusively performed in DIA mode. Orbitrap MS1 scans were recorded at a resolution of 240,000, a scan range from 380 to 980 m/z using 500% normalized AGC target and 100 ms maximum injection time. The window width of the Astral MS/MS scans was optimized in accordance to the sample’s precursor density. A total of 60 variable width isolation windows were defined which covered the precursor selection range of 380 to 980 m/z (see **Extended Data Table 2**). Fragment ion spectra were acquired with 20 ms maximum injection time, 500% normalized AGC target, and 25% HCD collision energy.

### Raw data processing

All 792 raw files were analyzed in a combined library-free search using DIA-NN (version 1.8.1)^34^. A human reference proteome FASTA file from UniProt (UP000005640, including reviewed entries, downloaded March 22, 2024) was provided. The search was performed with match-between-runs enabled and mass accuracy set to 8 and MS1 accuracy to 4 and the scan window radius to 6. Protein inference was based on genes using heuristic protein grouping. A maximum of one missed cleavage was allowed. The output “report.tsv” file was normalized using directLFQ (v.0.2.19) at standard settings^35^. The resulting ‘report.tsv.protein_intensities.tsv’ file was used for downstream data analysis steps. No data imputation was applied.

### Bioinformatics data analysis

#### i) Data filtering and quality control

Initial data processing was performed using Python (v3.12.5). At the sample level, measurements were excluded where the number of identified proteins was either 1.5 standard deviations below or 3 standard deviations above the median number of identifications across all samples. Quality control analyses, including assessment of protein group distributions and protein intensity rank plots, were performed on this filtered dataset to verify data quality and consistency. For subsequent analyses, an additional filtering criterion at the protein level was applied, requiring proteins to be detected in at least 70% of samples, if not stated otherwise. Where indicated for downstream analyses, data was standardized on the protein level using z-score normalization within each patient. By this, protein intensities are centered around the mean and are scaled to unit variance on a per-patient basis.

#### ii) Continuous analysis approach

To explore protein expression gradients along the zonation trajectories for each patient, a linear mixed model was fitted, using the portal score (*S*) as an independent variable, and protein expression (*y*) as the corresponding dependent outcome^25^. A random intercept effect on the patient the cell came from was added to control for differing expression baselines in different samples. Cells were removed by outlier intensity values using Tukey’s fences method^36^. The resulting expression intensities were normalized using a robust scaler (by which the median is removed and the inter-quartile range is divided). The full model is then given by following equation:

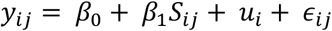

Where *y*_*ij*_ is the response variable for cell *j* in patient *i*, β_3_ is the fixed intercept, β_1_ is the fixed slope, from here on called zonation coefficient, for the predictor variable *S_ij_*, *u_i_* is the random intercept for patient *i*, where 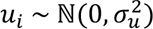, and *ε_ij_* is the residual error term, where 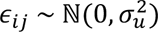

Once fitted, these models can be used to explore significant spatial expression gradients, by testing the zonation coefficient β_1_ against the null hypothesis that it equals zero (expression is spatially uniform in the central-portal axis) via a Wald test. The estimated zonation coefficients are compared to their standard error to determine the likelihood that the observed effect could occur by chance is determined. A protein expression gradient that increases from portal to central vein is indicated by a zonation coefficient significantly lower than zero. In contrast, an expression gradient that increases towards the portal vein is indicated by a significant positive zonation coefficient. The strength of the gradient is determined by the absolute value of the zonation coefficients (β_1_). Proteins with a zonation coefficient greater than 1 ((Ιβ_1_Ι > 1) and a q-value less than 0.05 were considered significantly zonated.

In order to evaluate whether protein zonation gradients differ between experimental conditions, the zonation coefficients (β_1_) of the linear mixed models fitted for controls were compared to those of subjects with fibrotic liver tissue. Only proteins with significant zonation gradients in the control condition were selected, and their zonation coefficients were compared using a Wald test. The null hypothesis that the difference in zonation coefficients (Δβ_1_) between the two conditions is zero is assessed by the Wald test. A test statistic that is distributed normally under the null hypothesis is obtained by dividing the observed difference by its standard error.

#### iii) Statistical Analysis and Data Visualization

Principal component analysis (PCA) was performed using scikit-learn after data standardization using StandardScaler, by which each feature is centered and scaled to unit variance. Prior to PCA, outliers were removed using a z-score based approach, where samples with any feature having an absolute z-score greater than 3 were excluded from further analysis. Explained variance ratios were calculated for each component. For spatial analysis, both continuous and discrete approaches were employed. The continuous analysis is explained in detailed below. For discrete analysis, data were divided into 20 equidistant bins along the portal-central axis, and mean protein expression was calculated for each bin. Z-scores were calculated using scipy.stats.zscore function along the bin axis, with NaN values omitted from the calculation. LOWESS smoothing (fraction=0.4) was applied to protein intensities plotted against the portal-central ratio of selected proteins to visualize their spatial behavior. Moreover, protein expression heatmaps were used for visualization and proteins were ordered by the absolute expression difference along the spatial trajectory. Hierarchical clustering was performed using Ward’s method with Euclidean distance to identify general protein expression patterns. Overrepresentation analysis was performed using Enrichr’s web interface with the complete dataset as background. Gene set enrichment analysis (GSEA) was performed using MSigDB collections (KEGG Pathways and MSigDB Hallmark). Statistical significance was assessed using Benjamini-Hochberg correction for multiple testing. For visualization of entire pathways, the median z score of the proteins assigned to the respective pathway were calculated. For comparison between analysis approaches, protein intensities were analyzed using both continuous (LMM-based) and discrete methods. For the discrete analysis, protein intensities were tested for spatial differences using one-way ANOVA implemented through ordinary least squares (OLS) regression with statsmodels. Expression differences between the 20 spatial bins were tested using Type II sum of squares. P-values were adjusted for multiple testing using the Benjamini-Hochberg procedure to control the false discovery rate. The statistical significance (-log10(q-value)) from both approaches was compared, and correlation between methods was assessed using Spearman correlation. A significance threshold of q < 0.05 was applied for both methods. Moreover, kernel density estimation with 20 density levels was used to visualize the distribution of significance values. Statistical comparisons between conditions were performed using two-sided Wilcoxon tests for paired samples and two-sided unpaired rank sum tests for absolute differences. Delta values for zonation coefficient changes were calculated as the median absolute difference between conditions. Regression analyses included 95% confidence interval calculations for trajectory visualization. First and third quartiles are shown by box plots with center line indicating median and whiskers extending to 1.5× interquartile range. All visualizations were generated using seaborn and matplotlib libraries in Python.

#### iv) Cross-species Comparison Analysis

Cross-species conservation of protein zonation was assessed by comparison of human liver data to the publicly available mouse scDVP dataset^15^. Protein homology mapping between human and mouse was performed using DIOPT (DRSC Integrative Ortholog Prediction Tool), by which multiple orthology prediction algorithms are integrated to provide comprehensive homology scores^37^. The protein filtering and normalization strategies described in the original paper were maintained. The mouse data was then re-analyzed using our continuous analysis approach to enable direct comparison of zonation patterns between species.

## Data availability

Extended Data Tables and generated code will be available upon publication.

---

**Extended Data Fig.1.**
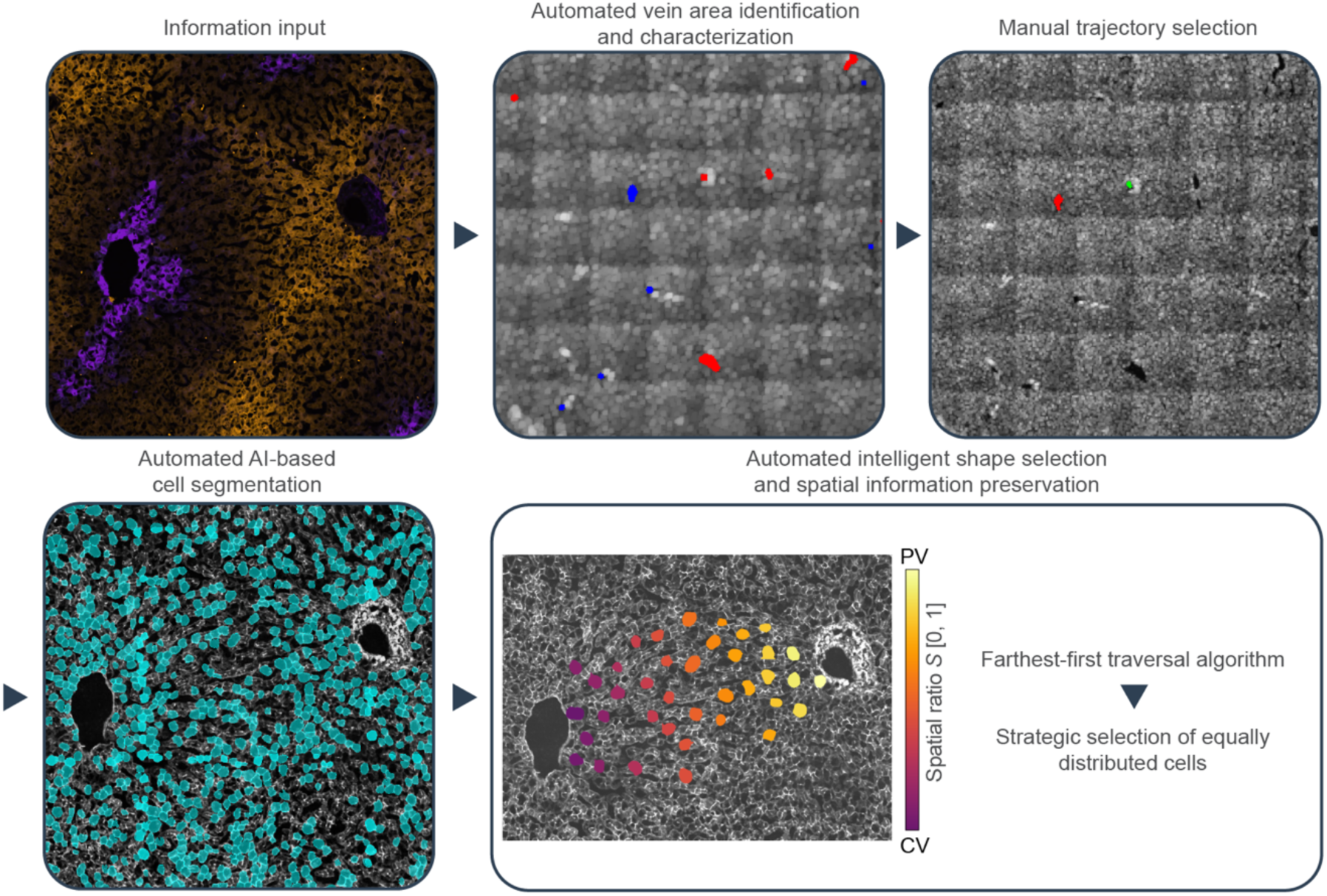
Automated selection of hepatocytes along zonation trajectories. The microscopy images are input for automated vein identification and characterization based on marker proteins. The user then manually selects suitable trajectories. Pre-segmented cell contours (turquois) are the basis for the farthest-first traversal algorithm, maximizing minimum distances between selected shapes to achieve uniform cell distribution along the portal-central axis. Selected shapes (colored dots) are assigned a portal score *S* from 0 (CV, central vein) to 1 (PV, portal vein) based on their relative position.

**Extended Data Fig.2.**
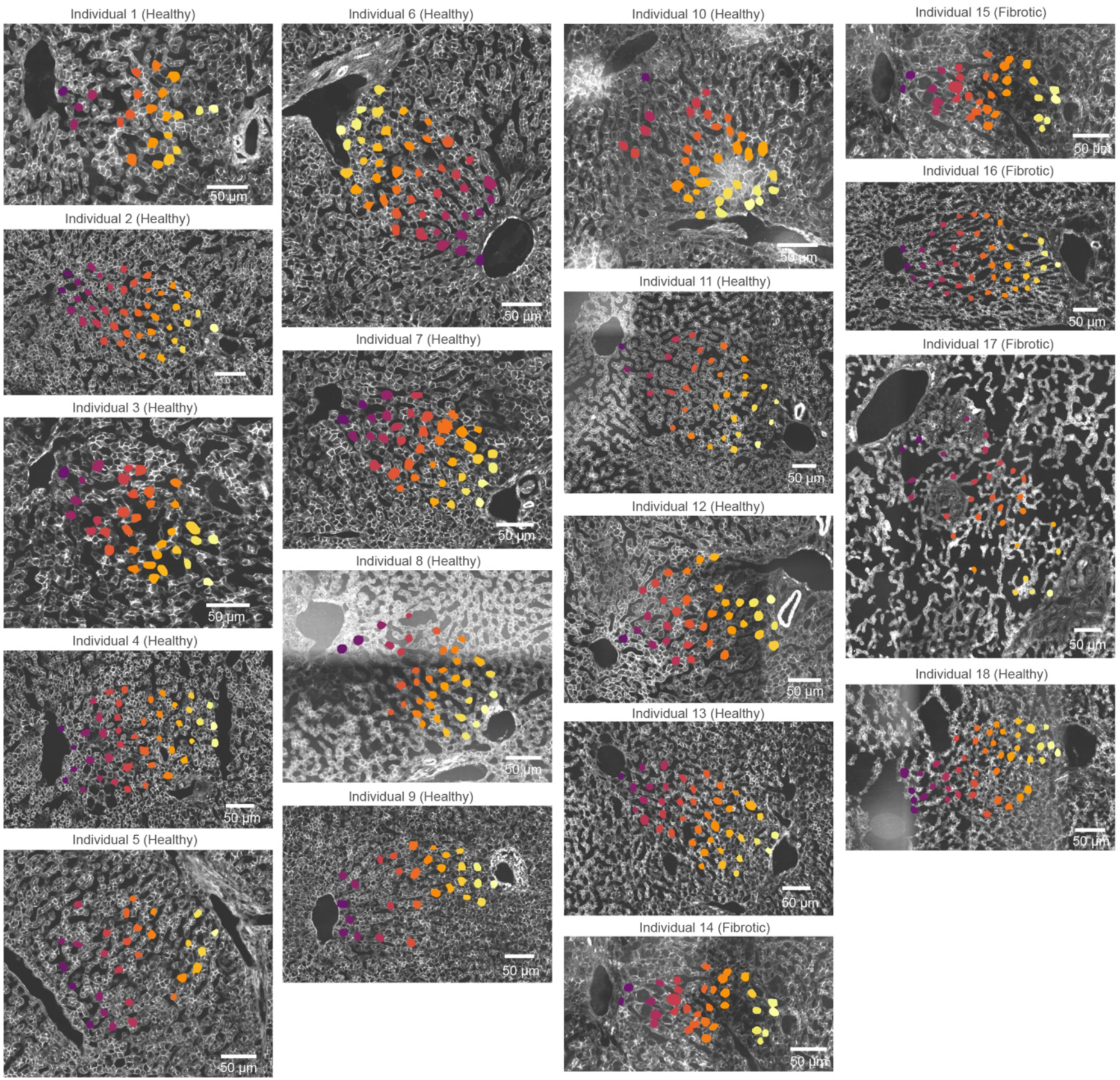
Overview of selected zonation trajectories across the human cohort. Representative microscopy images of liver sections from 18 individuals (healthy and fibrotic) highlighting hepatocytes along the chosen portal-central vein trajectories, which were subsequently isolated and measured by LC-MS (colored contours overlaid on phalloidin images). The color gradient indicates the spatial ratio *S* from a central vein (purple, *S* = 0) to the corresponding portal vein (yellow, *S* = 1). Scale bars, 50 μm.

**Extended Data Fig.3.**
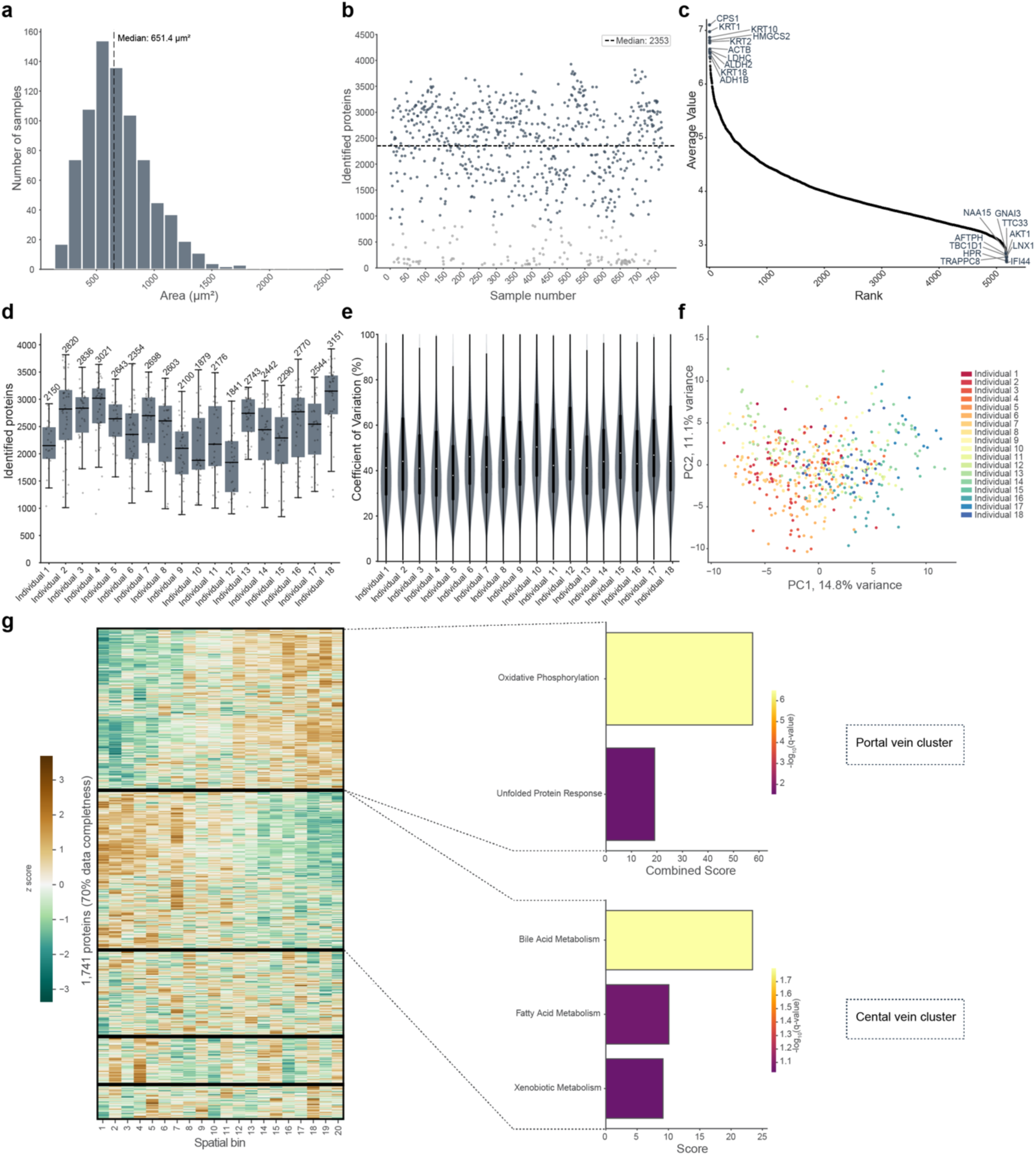
Data quality control of single-cell hepatic proteome dataset. a-c) Analysis of complete dataset before filtering (n = 792, N = 18): a) Histogram of all isolated hepatocyte shape areas. **b)** Number of identified proteins across all measured samples. The dotted line is the median number of quantified proteins. Grey points indicate samples with protein numbers below the threshold of 1.5 standard deviations from the median, which were excluded in later analysis steps (included = 649, excluded = 143). **c)** Protein abundance rank plot with the top and bottom 10 most abundant proteins highlighted. **d-f)** Analysis after standard deviation filtering per patient (n = 649, N = 18): **d)** Box plots showing protein identification numbers across all individuals with median values indicated above each box and proteins shown as grey dots with jitter. Box plots show first and third quartiles with a center line indicating the median and whiskers extend to 1.5× interquartile range. **e)** Violin and box plots showing coefficients of protein variation across all patients. **f)** Principal component analysis of the entire cohort after outlier removal with samples colored by the patient (n = 513, N = 18). **g)** Protein expression heatmap from healthy individuals across spatial bins using proteins detected in at least 70% of samples (n = 1741, N = 14). Proteins were clustered using Ward’s hierarchical clustering with Euclidean distance. Bar plots show overrepresentation analysis of the two largest clusters results using MSigDB Hallmark collection with the complete dataset as background and statistical significance assessed using Benjamini-Hochberg correction for multiple testing (q < 0.1). The clusters were classified as portal and central in accordance with the terms.

**Extended Data Fig.4.**
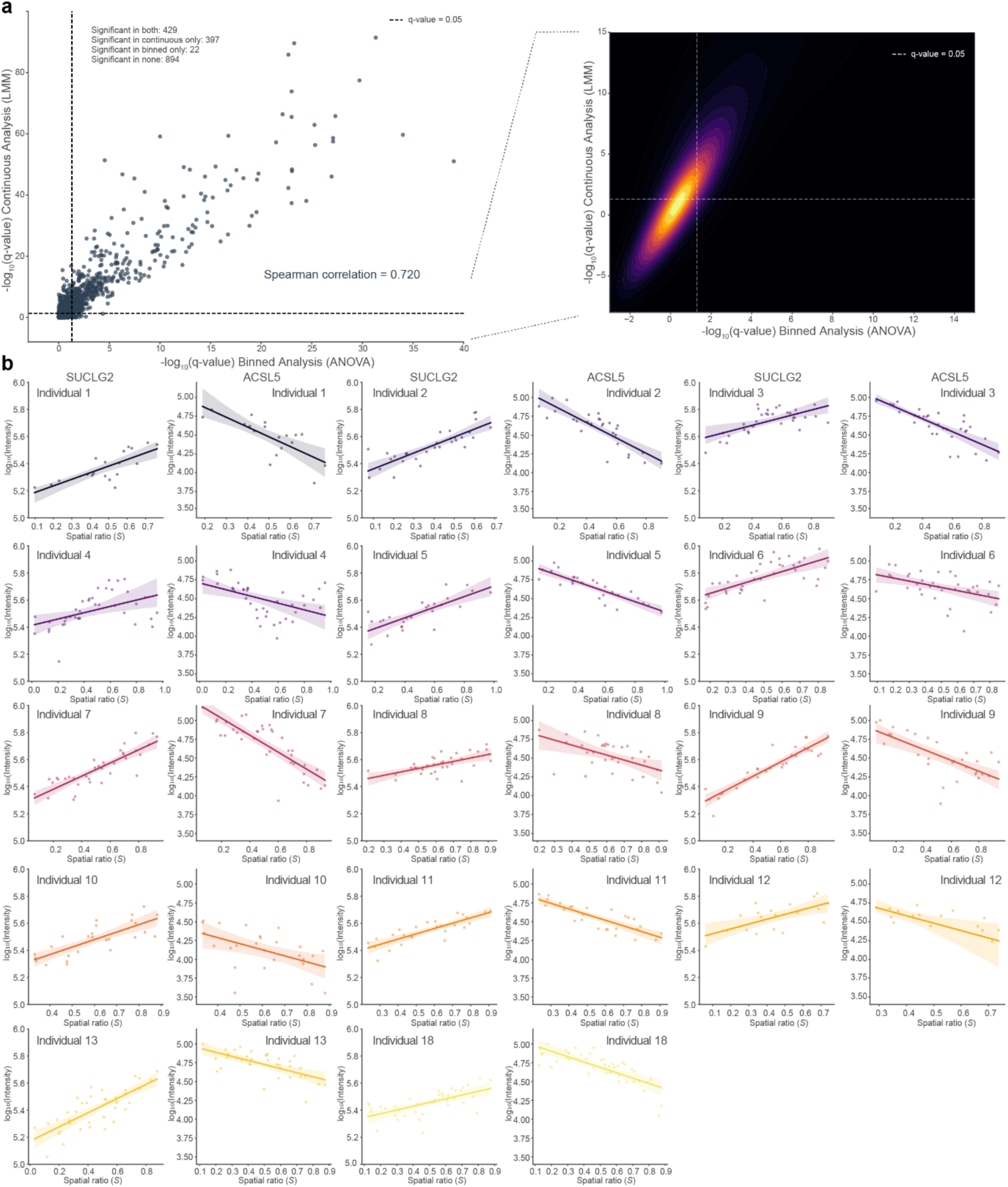
Individual patient analysis of the most significantly zonated proteins. **a)** Left: Statistical significance comparison between continuous (LMM-based) and binned (ANOVA-based) analysis. For the binned analysis, protein intensities were divided into 20 equal-width spatial bins and analyzed using one-way ANOVA to test for expression differences between bins. Each dot represents a protein with its respective -log10(q-value). Numbers of proteins significant in both, either, or neither analysis and Spearman correlation coefficient are indicated. Right: Two-dimensional kernel density plot showing the distribution of -log10(q-value) with 20 density levels in the enlarged region (x: -3, 15; y: -8, 15). Dashed lines indicate significance threshold (q < 0.05). **b)** Protein intensities (-log10 transformed) of the two most significantly zonated proteins (SUCLG2: periportal, ACSL5: pericentral) plotted against spatial ratio *S* per individual. Each dot represents a single hepatocyte measurement, solid lines show linear regression fits with 95% confidence intervals (shaded areas). Data include solely the healthy human cohort with proteins showing at least 70% data completeness (n = 1741, N = 14).

**Extended Data Fig.5.**
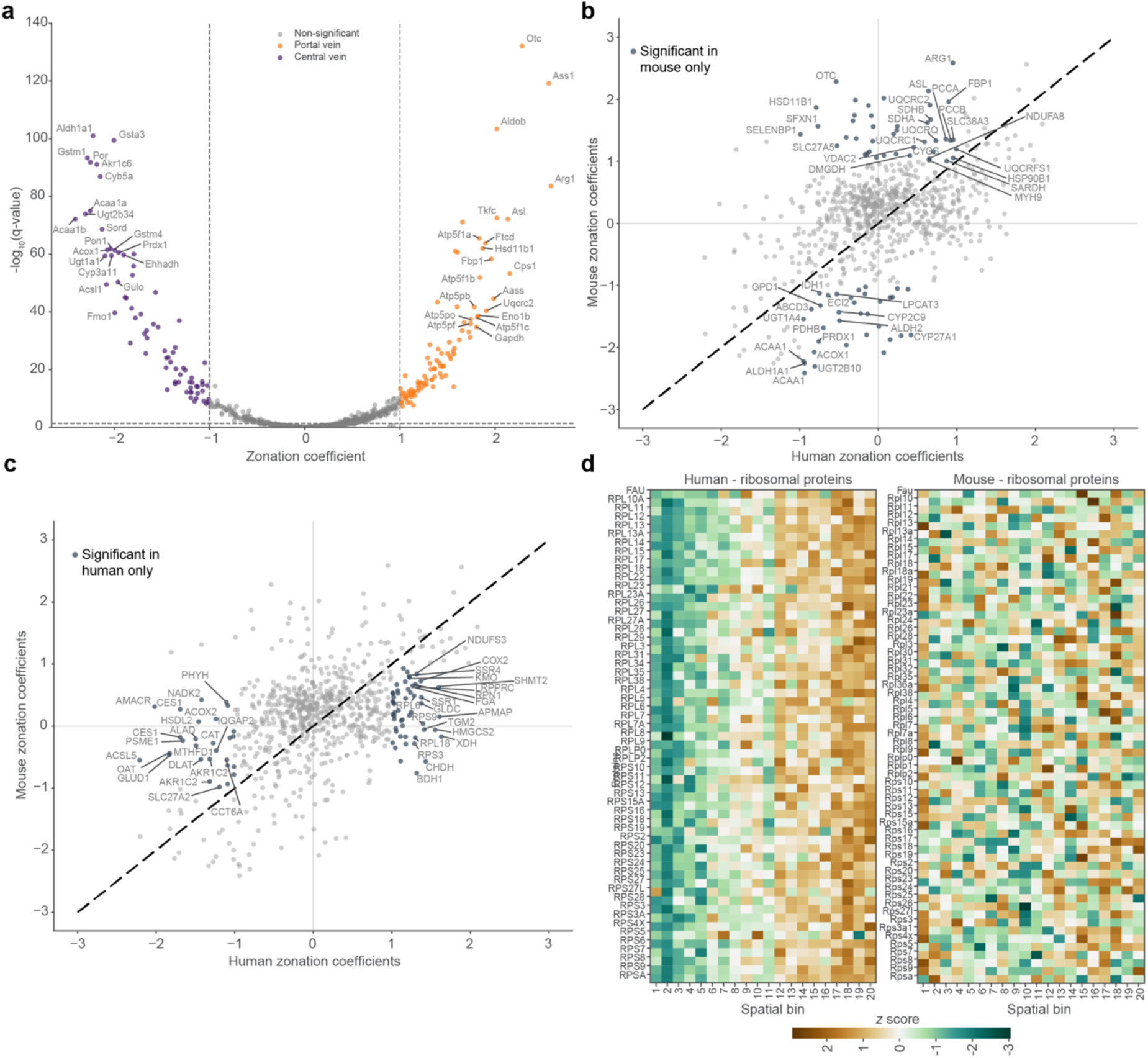
Cross-species analysis of liver zonation patterns. **a)** Application of continuous analysis to mouse scDVP data showing zonation coefficient β1 from linear mixed model plotted against -log10(q-value) after Wald test. Orange and purple dots indicate significantly enriched proteins (q < 0.05 and |coefficient| > 1) in the portal (PV; zonation coefficient > 1) and central vein (CV; zonation coefficient < -1) regions, respectively. **b)** Comparison of zonation coefficients between human and mouse shared proteins. Proteins with |coefficient| > 1 in mouse only are labeled and highlighted. **c)** Analysis as in b) highlighting proteins with |coefficient| > 1 in human liver tissue only. **d)** Protein expression heatmap (*z* scored) from detected proteins assigned to the KEGG ‘Ribosome’ term in humans (n=88) and mice (n=84). 20 equal-width spatial bins from central (spatial ratio *S* = 0) to portal (spatial ratio *S* = 1) are shown. Unless otherwise stated, only proteins at 70% data completeness are included (human: n = 1741, N = 14; mouse: n = 981, N = 3).

**Extended Data Fig.6.**
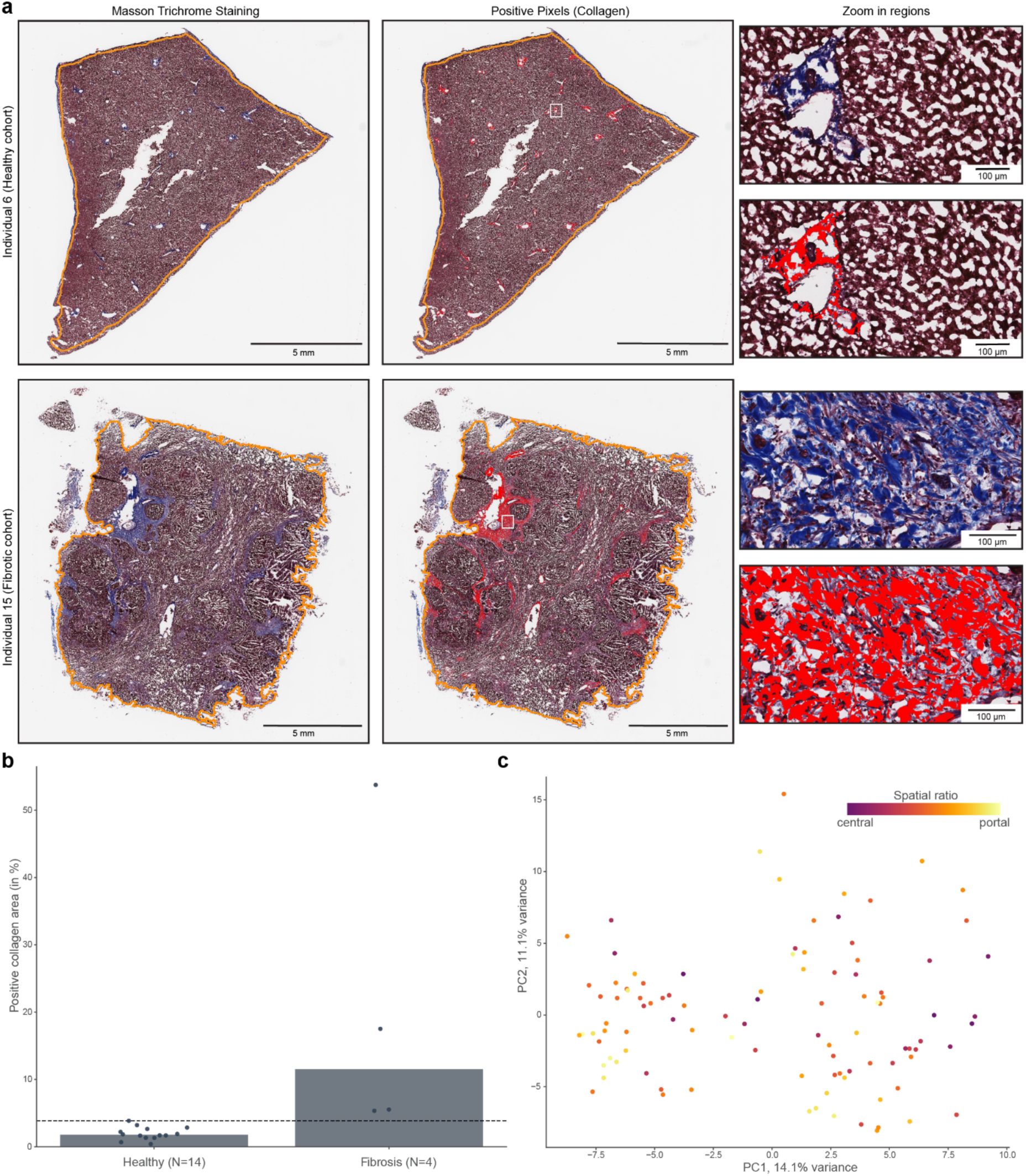
Loss of clear spatial architecture in fibrotic liver tissue. **a)** Masson Trichrome staining (left), corresponding collagen positive area detected by Area Quantification in HALO imaging software (middle) and zoom in regions (right). The tissue area considered in the analysis (excluding the liver capsule) is indicated by an orange border. Scale bars, 5mm and 100 μm. **b)** Percentage of positive collagen area on a tissue slide per individuum. The individuals are separated into healthy (N=14) and fibrotic (N=4). Dotted line represents the highest positive collagen area value detected in the healthy cohort. **c)** Principal component analysis of single-cell proteomes from fibrotic liver tissue. Each dot represents a single hepatocyte shape with color indicating its spatial ratio *S* (n = 100, N = 4).

## Notes

### Competing Interest Statement

M.M. is an indirect investor in Evosep. K.B. is co-founder and scientific advisor of Computomics GmbH, Tuebingen, Germany. All other authors declare no competing interests.

